# Gut and oral microbial community characterization from women with breast cancer, women with ductal carcinoma in situ, and healthy women reveals differences in gut but not oral microbiota

**DOI:** 10.1101/2024.07.22.604621

**Authors:** Emma McCune, Anukriti Sharma, Breanna Johnson, Tess O’Meara, Sarah Theiner, Maribel Campos, Diane Heditsian, Susie Brain, Jack A. Gilbert, Laura Esserman, Michael J. Campbell

## Abstract

This study characterized and compared the fecal and oral microbiota from women with early-stage breast cancer (BC), women with ductal carcinoma in situ (DCIS), and healthy women. Fecal and oral samples were collected from newly diagnosed patients prior to any therapy and characterized using 16S rRNA sequencing. Measures of gut microbial alpha diversity were significantly lower in the BC versus healthy cohort. Beta diversity differed significantly between the BC or DCIS and healthy groups and several differentially abundant taxa were identified. Clustering (non-negative matrix factorization) of the gut microbiota identified 5 bacterial guilds dominated by either *Prevotella*, Enterobacteriaceae, *Akkermansia*, Clostridiales, or *Bacteroides*. The *Bacteroides* and Enterobacteriaceae guilds were significantly more abundant in the BC cohort compared to healthy controls whereas the Clostridiales guild was more abundant in the healthy group. Finally, prediction of functional pathways identified 23 pathways that differed between the BC and healthy gut microbiota including lipopolysaccharide biosynthesis, glycan biosynthesis and metabolism, lipid metabolism, and sphingolipid metabolism. In contrast to the gut microbiomes, there were no significant differences in alpha or beta diversity in the oral microbiomes and very few differentially abundant taxa were observed. NMF analysis of the oral microbiota samples identified 7 guilds dominated by *Veillonella*, *Prevotella*, Gemellaceae, *Haemophilus*, *Neisseria*, *Propionibacterium*, and *Streptococcus,* however, none of these guilds were differentially associated with the different cohorts. Our results suggest that alterations in the gut microbiota, but not oral microbiota, may provide the basis for interventions targeting the gut microbiome to improve treatment outcomes and long-term prognosis.

**IMPORTANCE:** Emerging evidence suggests that the gut microbiota may play a role in breast cancer. Few studies have evaluated both the gut and oral microbiomes in women with breast cancer (BC) and none have characterized these microbiomes in women with ductal carcinoma in situ (DCIS). We surveyed the gut and oral microbiomes from women with BC or DCIS and healthy women and identified compositional and functional features of the gut microbiota that differed between these cohorts. In contrast, very few differential features were identified in the oral microbiota. These findings suggest that the oral microbiome is unlikely to be an effective risk marker for DCIS or breast cancer compared to the gut microbiome. Understanding the role of gut bacteria in BC and DCIS may open up new opportunities for the development of novel markers for early detection (or markers of susceptibility) as well as new strategies for prevention and/or treatment.

## INTRODUCTION

Breast cancer is a complex disease: its development has been linked to environmental, genetic, and biological risk factors, its progression can span decades, and its clinical course is quite variable.^1,2^ However, over half of the women who develop breast cancer have no known risk factors and few of the women with a genetic predisposition to breast cancer, or who have been exposed to known environmental risk factors, go on to develop the disease.^3,4^ Clearly, additional contributing factors need to be identified. One such factor that has gained recent attention is the human microbiome, the collection of commensal microbes living within and on the human body.^5,6^

Ductal carcinoma in situ (DCIS) is a heterogeneous proliferative condition with increased risk of developing invasive breast cancer. While only 15-45% of DCIS progresses to invasive cancer, this is still considered sufficient to warrant an aggressive surgical approach given the lack of criteria to distinguish indolent from aggressive disease. Here again, the human microbiome may play a role.

The gut microbiome harbors thousands of bacterial species which are affected by host genetics, lifestyle, and environmental factors. Alterations in these microbial communities have been linked to a variety of diseases including inflammatory bowel disease,^7,8^ obesity,^9^ diabetes,^10,11^ rheumatoid arthritis,^12^ cardiovascular diseases,^13^ and cancer.^14–20^ Gut microbes have been shown to influence local and systemic immunity and metabolism^20–23^ including systemic levels of estrogen and its metabolites.^16,18,24,25^ Comparisons of the gut microbiota of women with breast cancer and healthy women have revealed differences in the quantity and diversity of various genera.^26–31^ In addition, antibiotic use has been linked to increased risk of breast cancer, likely due to its effects on the gut microbiota.^32^

A link between periodontal disease and cancer has been suggested through several studies looking both at specific types of cancers and at the overall total cancer rate and the relationship to periodontal disease.^33^ The scientific rationale behind this association is that inflammation is a major factor in both periodontal disease and cancer. With respect to breast cancer, Hujoel and coworkers found a weak association between periodontal disease and breast cancer risk.^34^ Swedish researchers have reported a significant increased risk of breast cancer associated with chronic periodontal disease (odd ratio = 2.36).^35,36^ Although these results suggest a link between breast cancer the oral microbiota, two previous studies found no differences in microbial alpha or beta diversity and very few if any differentially abundant taxa in the oral microbiota from women with breast cancer compared to that from healthy women.^37,38^

Given the microbiome’s potential involvement in carcinogenesis, we hypothesized that changes in the oral and/or gut microbiota might be associated with the development of breast cancer. Moreover, there have been no studies evaluating the microbiomes of women with ductal carcinoma in situ (DCIS). Therefore, in this study we characterized the gut and oral microbiota of women with newly diagnosed invasive breast cancer (BC) or ductal carcinoma in situ (DCIS), and compared these to the gut and oral microbiota of healthy women recruited from an age-matched population.

## RESULTS

### Participant characteristics

A total of 185 women were enrolled and received home sample collection kits for this study. This included 85 women with breast cancer (BC), 40 women with ductal carcinoma in situ (DCIS), and 60 healthy women (**Supplementary Figure 1A**). Eighty-three percent (n=154) of the sample kits were returned (73 BC, 32 DCIS, 49 healthy). As shown in **Table 1**, women in each of the three groups were similar for age, body mass index (BMI), and breast density.

**Table 1.**
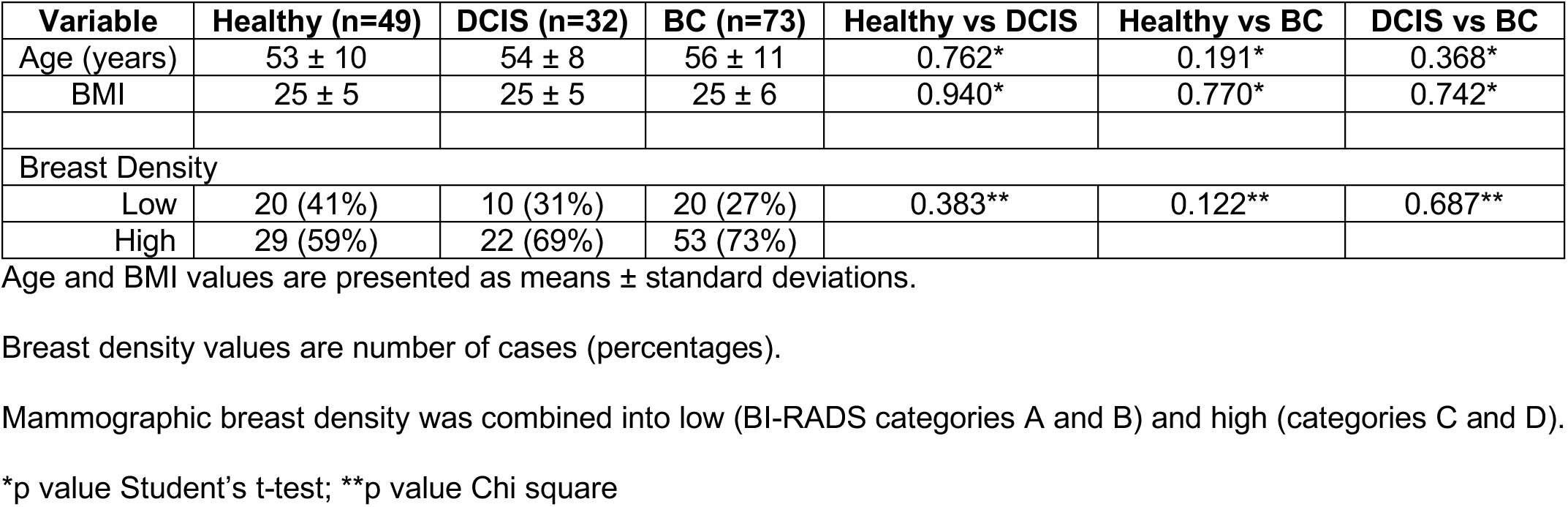
Demographics of study participants.

Approximately 40% of oral samples failed to yield sequencing results, typically due to low DNA yield, whereas only 10% of the gut samples failed to yield sequencing results. Overall we analyzed 137 gut samples (66 from women with BC, 29 from women with DCIS, and 42 from healthy women) and 93 oral samples (47 from women with BC, 15 from women with DCIS, 31 from healthy women)(**Supplementary Figure 1A**). Paired gut and oral microbiome data from the same individual were obtained from 40 BC, 12 DCIS, and 24 healthy participants (**Supplementary Figure 1B**).

### Variations in gut and oral microbiota diversity in women with breast cancer, women with DCIS, and healthy women

The microbial α-diversity of gut and oral samples was compared between cohorts based on Shannon and Simpson diversity indices. As shown in **Figure 1A**, the gut microbiota of women with BC had a statistically significant lower α-diversity compared to healthy women as indicated by the Shannon index (p=0.049) and the Simpson index (p=0.011). No differences in gut microbial α-diversity were observed between women with DCIS and women with BC or healthy women. The Shannon and Simpson indices for the oral microbiome did not differ across the three cohorts (**Figure 1C**).

**Figure 1.**
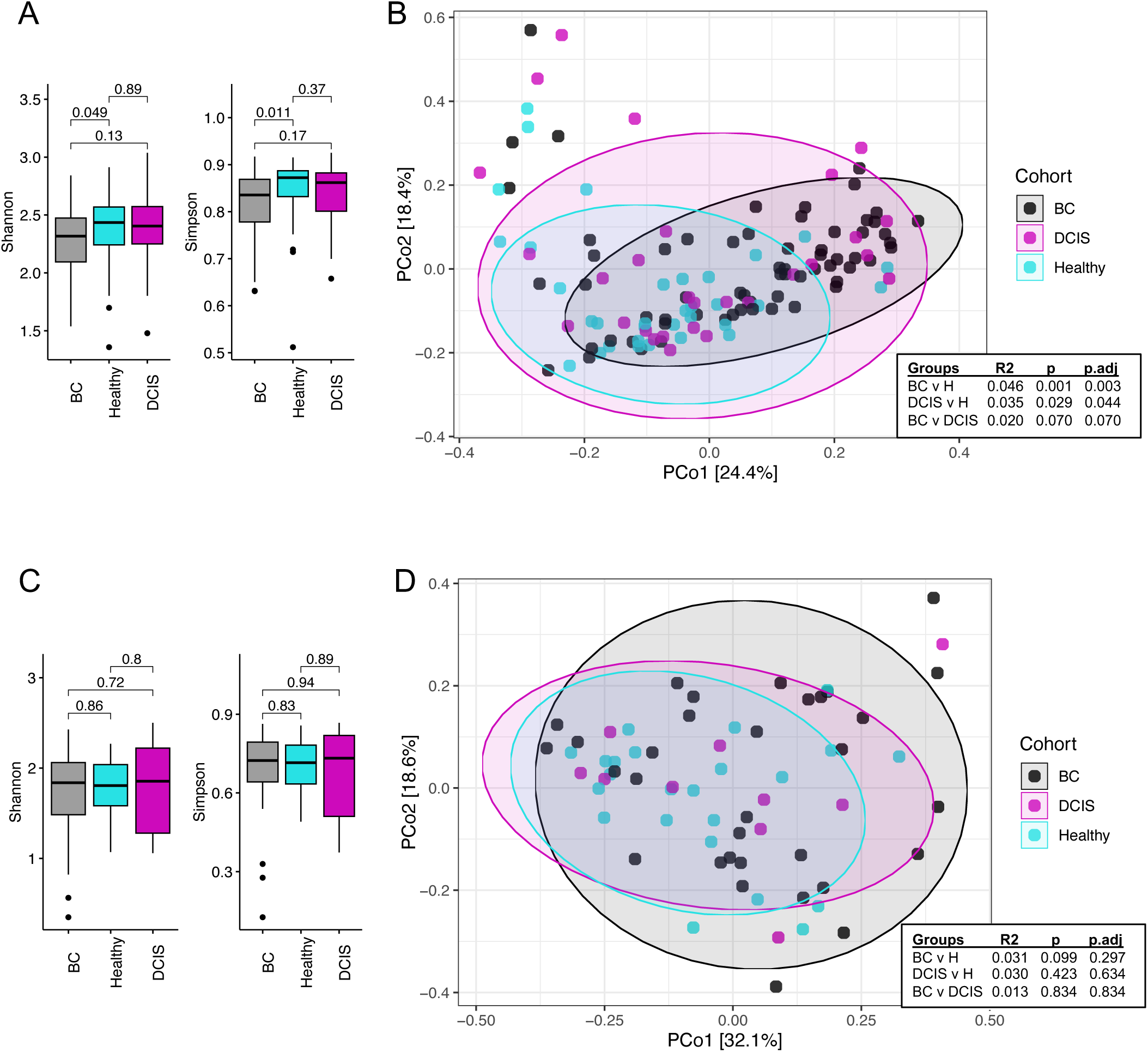
Diversity analysis of the oral and gut microbiota among the three cohorts. Shannon and Simpson alpha-diversity analyses at the genus level are shown for gut (A) and oral (C) samples. p-values calculated using Wilcoxon tests. Principal-coordinate analysis (PCoA) based on Bray-Curtis distance at the genus level are shown for gut (B) and oral (D) samples. R2 and p-values were calculated using pairwise PERMANOVA tests.

Differences in β-diversity were evaluated using principal-coordinate analyses (PCoA) based on Bray-Curtis distance metrics and pairwise PERMANOVA tests. Gut microbial β-diversity was significantly different between BC and healthy groups (p=0.001), as well as between the DCIS and healthy groups (p=0.032)(**Figure 1B**). In contrast, no differences in oral microbial β-diversity were observed between any group (**Figure 1D**).

### Comparison of gut and oral microbiota composition between women with breast cancer, women with DCIS, and healthy women

Taxa (phyla, families, genera) with a relative abundance ≥ 1% in the gut and oral microbiota in each cohort are presented in **Supplementary Fig 2**. Firmicutes and Bacteroidetes were the two most abundant phyla (together accounting for ∼90%) in the gut microbiota of all three cohorts (**Supplementary Figure 2A**). The fecal Firmicutes to Bacteroidetes ratio (F/B ratio) in the healthy group was significantly higher than in the BC group (p=0.027; **Supplementary Figure 3A**) but similar to that in the DCIS group (p=0.55). In the oral microbiota, Firmicutes, Proteobacteria, and Bacteroidetes were the three most abundant phyla (together accounting for 80-90%) across all three cohorts (**Supplementary Figure 2B**). In contrast to the fecal F/B ratios, the oral F/B ratios were similar in all groups (**Supplementary Figure 3B**).

Seventeen families were identified in the gut microbiota with a relative abundance ≥ 1% (**Supplementary Figure 2C**). Lachnospiraceae, Bacteroidaceae, and Ruminococcaceae together represented ∼70% of the relative abundance of family taxa. In the oral microbiota, 13 families were identified with a relative abundance ≥ 1%, with Streptococcaceae representing ∼40-50% and Pasteurellaceae, Veillonellaceae, Gemellaceae, Prevotellaceae, & Neisseriaceae representing another 30-40% (**Supplementary Figure 2D**).

At the genus level in the gut microbiota, 17 genera were identified with relative abundance ≥ 1% (**Supplementary Figure 2E**). *Bacteroides* was the most prevalent in all groups (BC: 28%, DCIS: 23%, Healthy: 20%) followed by *Blautia* (BC: 7%, DCIS: 6%, Healthy: 8%), *Faecalibacterium* (BC: 5%, DCIS: 4%, Healthy: 6%), *Ruminococcus* (BC: 5%, DCIS: 5%, Healthy: 4%), *Prevotella* (BC: 3%, DCIS: 5%, Healthy: 7%), *Akkermansia* (BC: 3%, DCIS: 3%, Healthy: 3%), and *Coprococcus* (BC: 2%, DCIS: 3%, Healthy: 3%). The remaining genera each represented ≤ 2% of the relative abundance in each cohort. Of the 13 genera identified with relative abundance ≥ 1% in the oral microbiota (**Supplementary Figure 2F**), *Streptococcus* was the most prevalent genus in all groups (BC: 42%, DCIS: 54%, Healthy: 48%) followed by *Veillonella* (BC: 11%, DCIS: 9%, Healthy: 9%), *Haemophilus* (BC: 10%, DCIS: 6%, Healthy: 8%), *Prevotella* (BC: 7%, DCIS: 7%, Healthy: 5%), *Neisseria* (BC: 5%, DCIS: 3%, Healthy: 5%), *Gemella* (BC: 2%, DCIS: 1%, Healthy: 3%), and *Rothia* (BC: 1%, DCIS: 3%, Healthy: 1%). The remaining genera each represented ≤ 2% of the relative abundance in each cohort.

### Identification of microbial taxa that differentiate DCIS, IDC, and healthy oral and gut microbiota

We used LEfSe analysis (with an LDA score cut off ≥ 3) to identify key taxa responsible for the differences in the compositions of the oral and fecal microbiotas between the three groups (**Figure 2**). In the fecal microbiota, 6 taxa were more abundant in the BC group compared to the healthy group including the family Bacteroidaceae and its corresponding genus *Bacteroides*; the class Actinobacteria; the family [Tissierellaceae] and corresponding genus *Finegoldia*; and the phylum Tenericutes (**Figure 2A**). The phylum Firmicutes and its corresponding class Clostridia, order Clostridiales, family Lachnospiraceae, and genera *Coprococcus* and *Anaerostipes* were more abundant in the healthy group compared to the BC group (**Figure 2A**). Comparing the fecal microbiota of the healthy and DCIS groups, 9 taxa were more abundant in the DCIS group (**Figure 2B**). These included the phylum Actinobacteria and its corresponding class Actinobacteria, order Actinomycetales, family Actinomycetaceae, and genus *Varibaculum;* the family Corynebacteriaceae and corresponding genus *Corynebacterium*; and the genera *Dialister* and *Megamonas.* Two taxa, the family Lachnospiraceae and corresponding genus *Faecalibacterium*, were more abundant in the healthy group compared to the DCIS group (**Figure 2B**). Finally, only 4 taxa were differentially abundant in the BC vs DCIS groups (**Figure 2C**). The genera *Phascolarctobacterium* and *Megamonas* were enriched in the BC gut microbiota whereas the family Actinomycetaceae and genus *Dialister* were enriched in the DCIS group.

**Figure 2.**
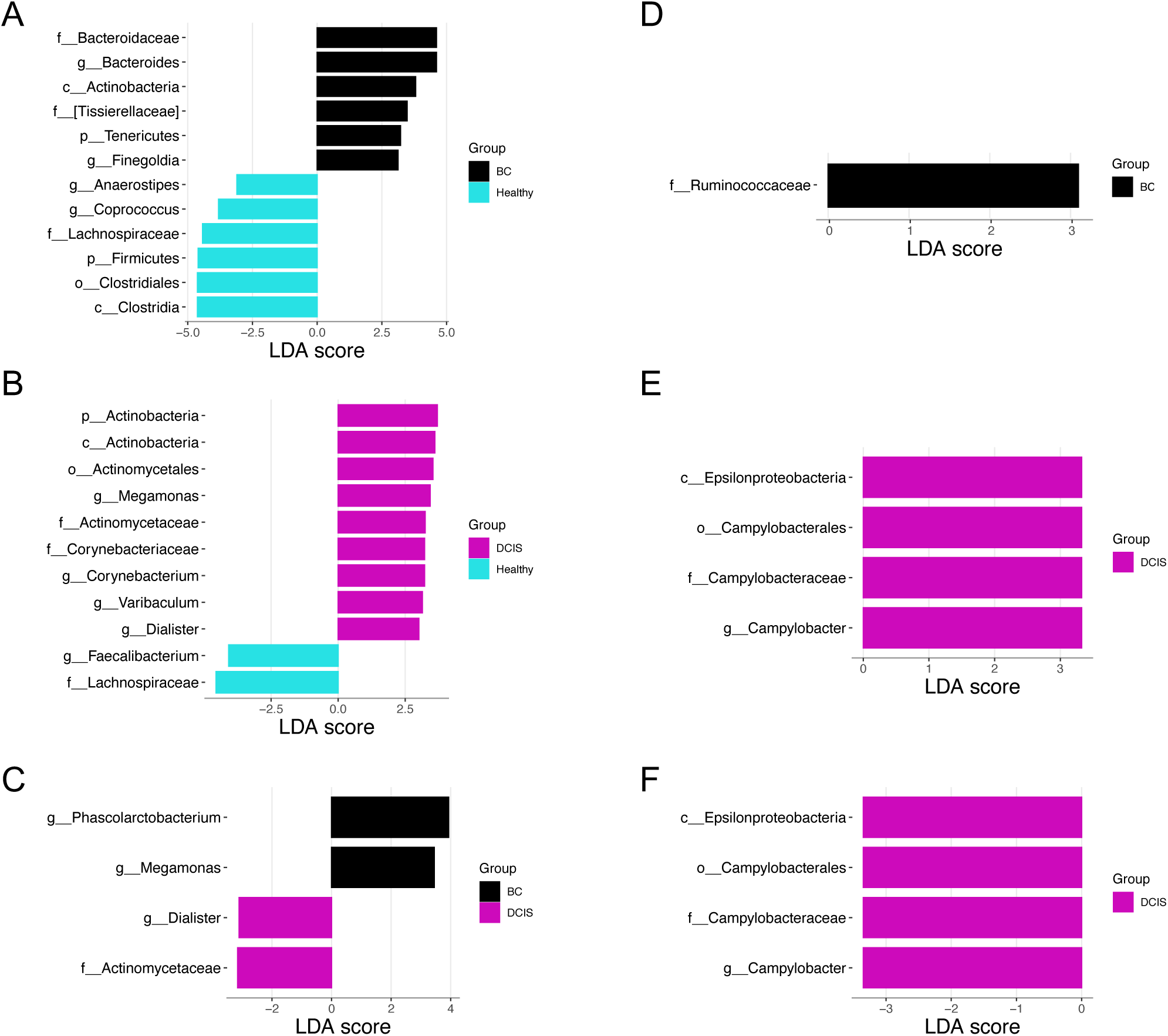
Taxonomic biomarkers identified by LEfSe. LDA bar graphs plotted from LEfSe analyses of gut (A-C) and oral (D-F) microbiota. The color of the bars represent the cohort. The logarithmic LDA score threshold was 3.0 for discriminative features. (A and D) BC vs healthy; (B and E) DCIS vs healthy; (C and F) BC vs DCIS.

In contrast to the gut microbiota, very few differentially abundant taxa were identified in the oral microbiota. The family Ruminococcaceae was enriched in the BC group compared to the healthy cohort (**Figure 2D**). The class Epsilonproteobacteria and corresponding order Campylobacterales, family Campylobacteraceae, and genus *Campylobacter* were more abundant in the DCIS gut microbiota compared to the BC and the healthy groups (**Figure 2E & 2F**).

DESeq2 analyses were also used to identify differentially abundant taxa at the genus level. As shown in **Figure 3A**, comparing the gut microbiota of the healthy cohort with the BC cohort, the genera *Coprobacillus*, *Parabacteroides*, *Streptococcus*, *WAL*, *Corynebacterium*, *Anaerococcus*, *Bacteroides*, *Acidaminococcus*, *Eggerthella*, *Peptoniphilus*, and *Finegoldia* were enriched in the BC cohort compared to the healthy cohort, while *Anaerostipes* was enriched in the healthy cohort.

**Figure 3.**
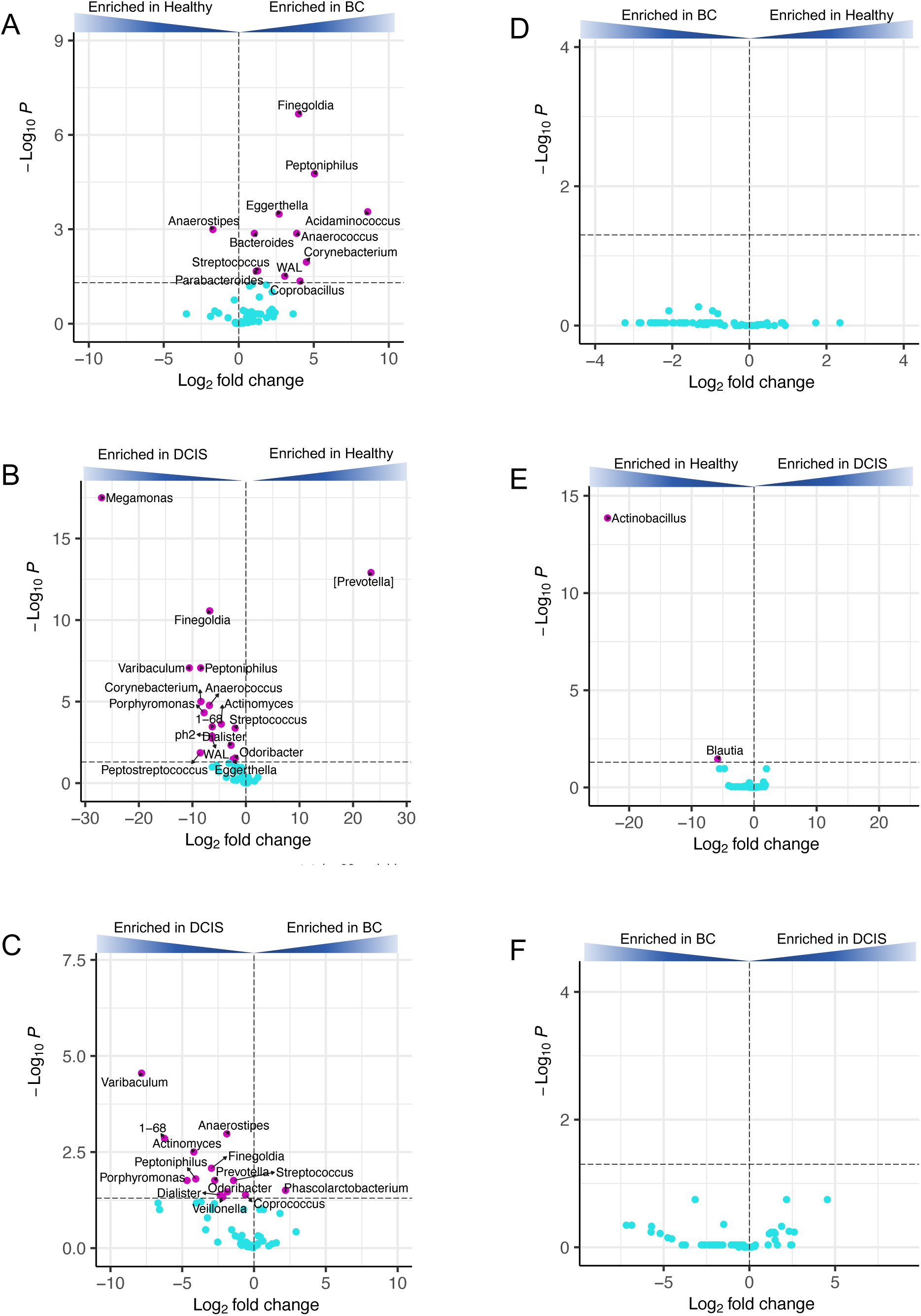
DESeq2 differential abundance analysis. Volcano plots of log2 fold differences in genera abundance in the gut microbiota (A, B, C) or oral microbiota (D, E, F) between BC and healthy (A, D), DCIS and healthy (B, E), and BC and DCIS (C, F). Dashed horizontal lines reflect adjusted p-value = 0.05. Names of differentially abundant genera (p.adj > 0.05) are shown (magenta dots). Blue dots represent not significant genera.

Comparing the healthy cohort with the DCIS cohort (**Figure 3B**) DESeq2 analysis identified 16 genera enriched in the DCIS cohort including 7 genera that were also differentially enriched in BC vs healthy (*Finegoldia*, *Peptoniphilus*, *Eggerthelia*, *Anaerococcus*, *Corynebacterium*, *WAL*, and *Streptococcus*). Only 1 genus, [*Prevotella*], was enriched in the healthy cohort compared to the DCIS group. Finally, comparing the DCIS cohort with the BC cohort, 13 genera were more abundant in the gut microbiota of the DCIS cohort compared to the BC cohort and only 1, *Phascolarctobacterium*, was enriched in the BC group (**Figure 3C**).

In contrast to the gut microbiota, very few differentially abundant genera were identified in the oral microbiota by DESeq2 analysis. No genera were found to be significantly enriched when comparing the BC and healthy groups (**Figure 3D**) or comparing the BC and DCIS groups (**Figure 3F**). Comparing the DCIS and healthy oral microbiota, only 2 genera, *Actinobacillus* and *Blautia*, were found to be differentially enriched (higher in healthy vs. DCIS)(**Figure 3E**).

### Gut and oral microbial guilds associated with breast cancer

Since community structure may be more informative than abundance differences of individual taxa, we clustered the gut and oral microbiota into guilds using non-negative matrix factorization (NMF). We identified 5 guilds in the gut microbiota samples and named them according to the most notable taxa: G1-*Prevotella*, G2-Enterobacteriaceae, G3-*Akkermansia*, G4-Clostridiales, and G5-*Bacteroides* (**Figure 4A**). The G4-Clostridiales guild is also characterized by higher abundances of the related families Ruminococcaceae and Lachnospiraceae and the related genera *Blautia*, *Faecalibacterium*, and *Ruminococcus*. The G2-Enterobacteriaceae and G5-*Bacteroides* guilds were significantly more abundant in the gut microbiota of women with breast cancer compared to healthy women, whereas the G4-Clostridiales guild was significantly more abundant in the healthy cohort compared to the BC group (**Figure 4B**).

**Figure 4.**
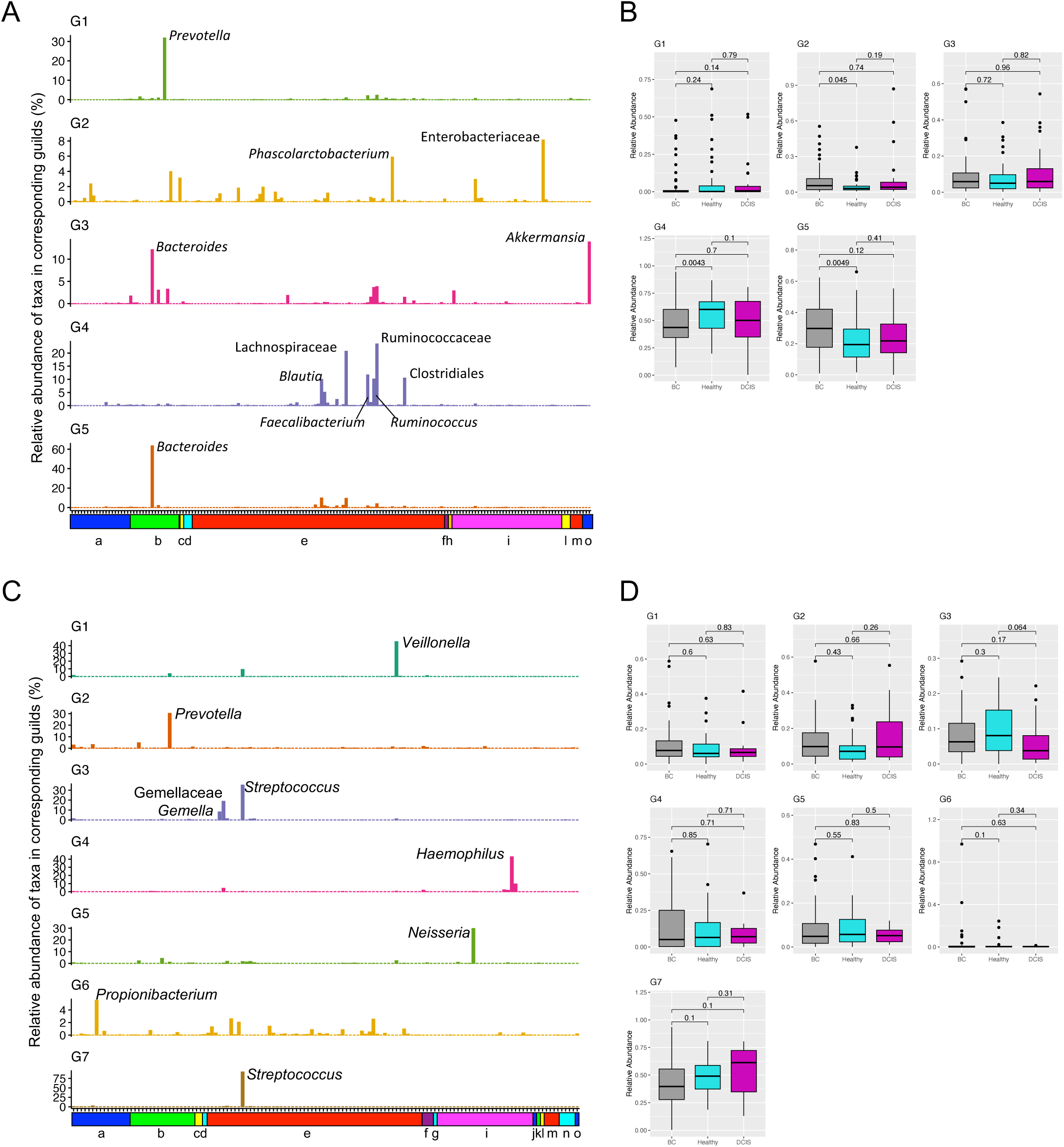
Microbiota guilds identified by non-negative matrix factorization analyses. (A and C) Each bar represents the relative abundance of a taxa in the guild (A: gut; C: oral). Color annotation bars indicates phyla for each taxa: (a) Actinobacteria, (b) Bacteroidetes, (c) Cyanobacteria, (d) Euryarchaeota, (e) Firmicutes, (f) Fusobacteria, (g) GN02, (h) Lentisphaerae, (i) Proteobacteria, (j) Spirochaetes, (k) SR1, (l) Synergistetes, (m) Tenericutes, (n) TM7, (o) Verrucomicrobia. (B and D) Box plots showing relative abundance of each guild by cohort (B: gut; D: oral). p-values calculated using Wilcoxon tests.

NMF applied to the oral microbiota samples identified 7 guilds which were named according to the most notable taxa: G1-*Veillonella*, G2-*Prevotella*, G3-Gemellaceae, G4-*Haemophilus*, G5-*Neisseria*, G6-*Propionibacterium*, and G7-*Streptococcus* (**Figure 4C**). In contrast to the gut guilds, and consistent with the LEfSe and DESeq2 analyses of the oral microbiota, none of these guilds were differentially abundant across the cohorts (**Figure 4D**).

### Predicted functional potential of the oral and gut microbial communities

While the taxonomic classifications of the bacterial communities of the three cohorts is useful, we were also interested in the functional potential of the oral and gut microbiota and how this might vary across the cohorts. Different methods are available that use 16S rRNA sequence data to infer the microbial genomes present and their potential functionality in the absence of whole-genome sequencing data.^39–41^ It should be noted that these methods are based on inference from known genomes and may not fully recapitulate the existing metagenomic content.

We performed a predictive functional analysis on the gut 16S sequence data using the Tax4Fun algorithm and LEfSe analysis to identify pathways differentially present in the three cohorts (**Figure 5**). Twenty-three pathways differed between the BC and healthy cohorts, 14 associated with the BC group and 9 with the healthy group (**Figure 5A**). Modules enriched in the BC group included lipopolysaccharide biosynthesis, glycan biosynthesis and metabolism, lipid metabolism, galactose metabolism, and sphingolipid metabolism. Modules enriched in the healthy controls included membrane transport, ABC transporters, cell motility, bacterial chemotaxis, flagellar assembly, and arginine and proline metabolism. Only 3 pathways differed between the DCIS and healthy cohorts, all three being enriched in the DCIS samples (**Figure 5B**). Finally, 16 pathways were identified as differentially present in the BC and DCIS cohorts, 8 enriched in the BC group and 8 enriched in the DCIS group (**Figure 5C**). Interestingly, most of the modules that differentiated BC and DCIS also differentiated the BC and healthy groups.

**Figure 5.**
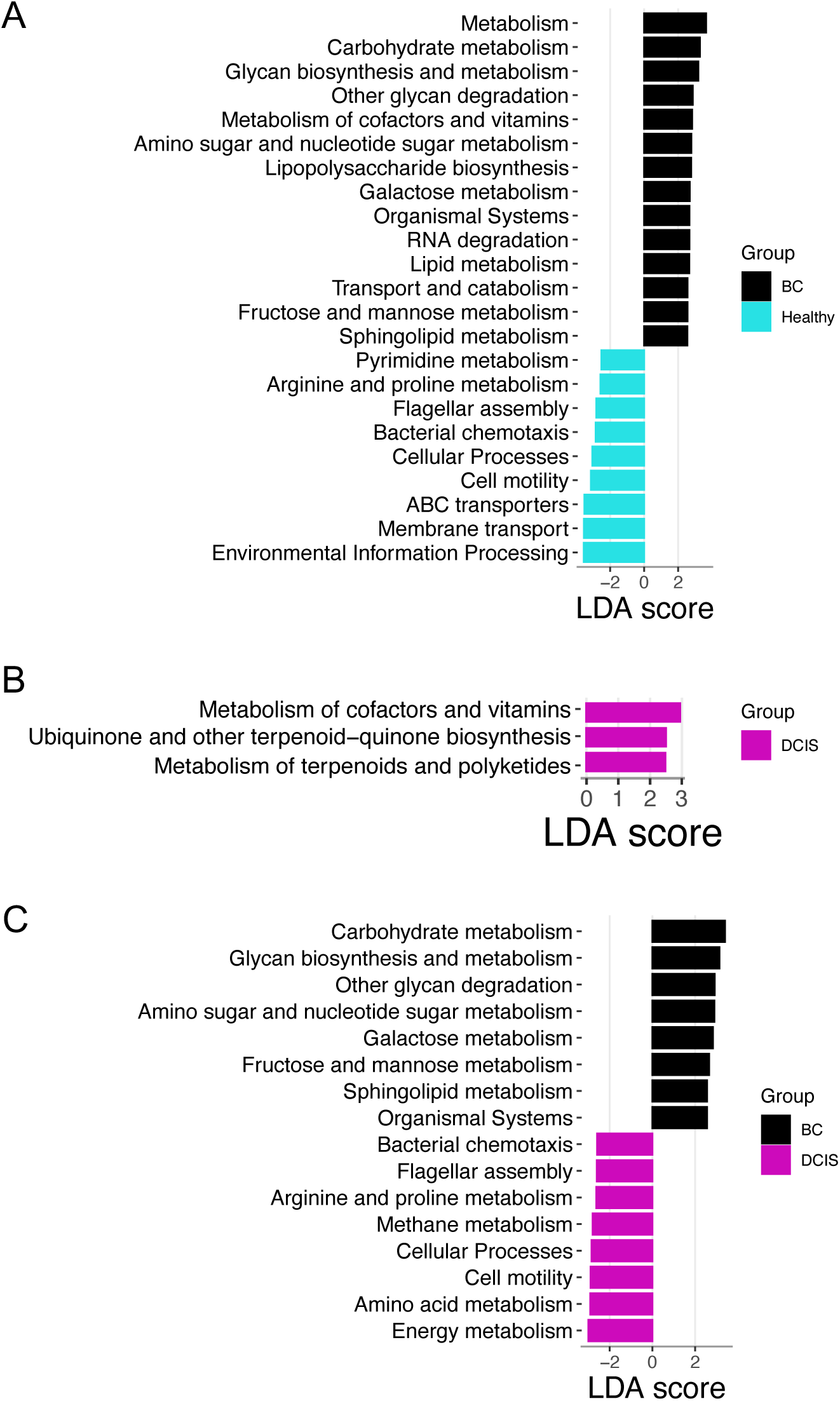
Functional analysis prediction. Tax4Fun was used to predict functional potential of microbiota using 16S rRNA gene sequence data. Differentially enriched bacterial functions among groups were identified using LEfSe analysis (logarithmic LDA score threshold = 2.5; p<0.05). The color of the bars represent the cohort. (A) BC vs healthy; (B) DCIS vs healthy; (C) BC vs DCIS.

In contrast to the gut microbiota, predictive functional analysis of the oral 16S sequence data demonstrated a lack of pathways discriminating the cohorts. No pathways were found to differentiate the BC and healthy or the DCIS and healthy cohorts and only 1 pathway (amino acid metabolism) was found to be enriched in the BC oral microbiota compared to the DCIS group (data not shown).

### Associations of gut and oral microbiota with breast density

Mammographic breast density, categorized as high (BI-RADS A and B) or low (BI-RADS C and D), did not differ significantly across the three cohorts (**Table 1**). Alpha diversity of the fecal microbiota and the oral microbiota did not differ between women with high vs. low mammographic density (**Figure 6A & 6B**). There were no differences in the F/B ratios between the high and low breast density groups in either the gut or oral microbiota. LEfSe analysis identified 7 taxa (LDA score ≥ 3) with differential abundance in the gut microbiota (**Figure 6C**). Six taxa were more abundant in the high breast density group, including the family Bacteroidaceae and corresponding genus *Bacteroides*; the phylum Cyanobacteria and corresponding class 4C0d-2 and order YS2; and the genus *Phascolarctobacterium*. The family Christensenellaceae was enriched in the low density group. No taxa were differentially abundant in the oral microbiota when comparing high vs low breast density groups.

**Figure 6.**
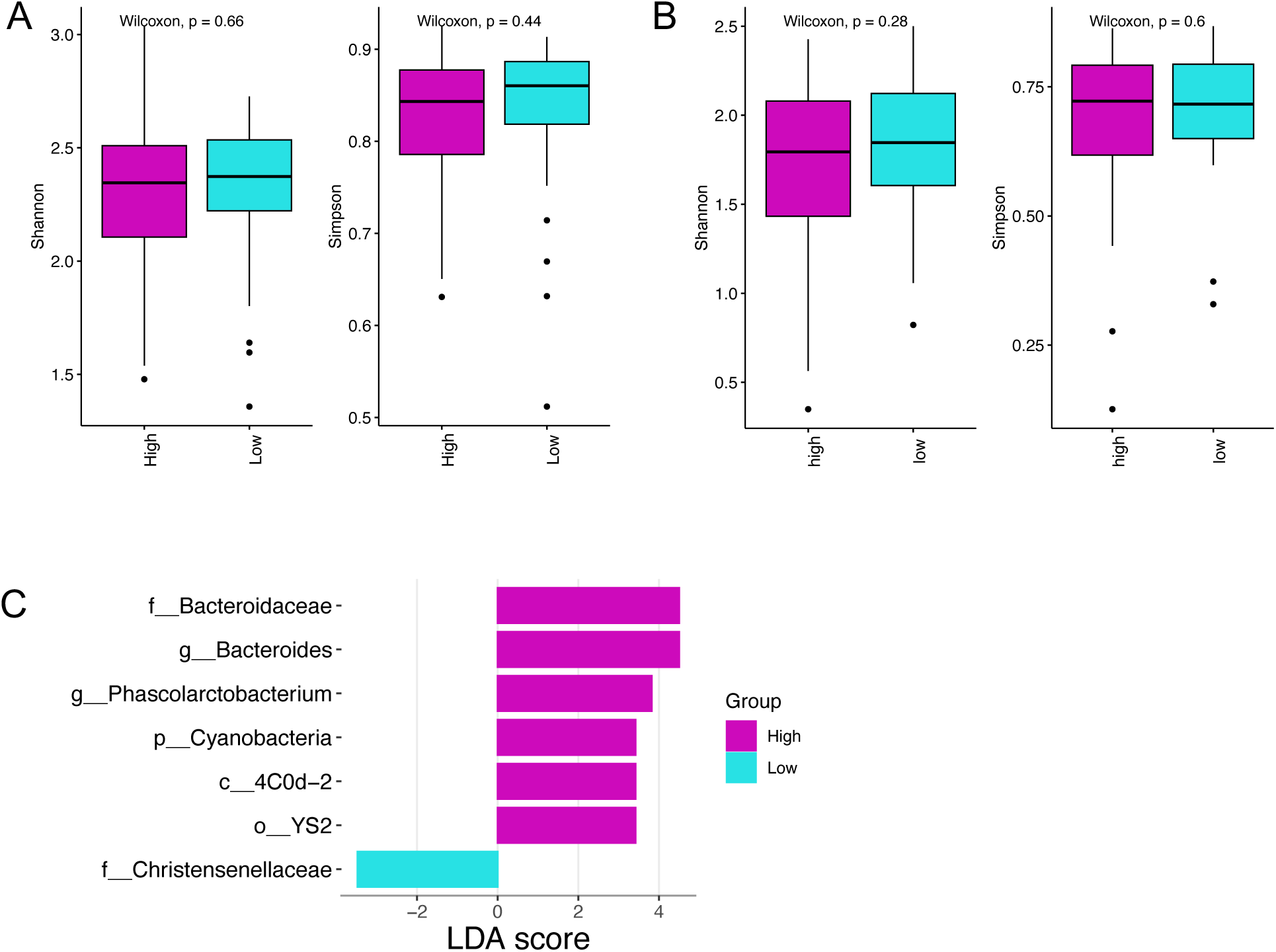
Oral and fecal microbiota features associated with mammographic breast density. Shannon and Simpson alpha-diversity analyses are shown for gut (A) and oral (B) samples comparing high and low breast density. (C) Differentially enriched taxa abundance in fecal samples between high and low breast density were identified using LEfSe analysis (logarithmic LDA score threshold = 3; p<0.05).

### Overlapping taxa between the oral and gut microbiotas

Because we characterized both gut and oral microbiota samples, we explored the similarity between these microbiotas. As shown in **Supplementary Fig4A**, the gut microbiota demonstrated significantly greater α-diversity compared to the oral microbiota. β-diversity, evaluated using principal-coordinate analyses (PCoA) based on Bray-Curtis distance metrics, significantly separated the gut and oral microbiota (**Supplementary Fig4B**).

LEfSe analysis was used to identified differentially abundant taxa between the gut and oral microbiomes. **Supplementary Figure 5** shows the top 10 (by LDA score) differentially abundant taxa at the phylum, family, genus, and species levels. At the phylum level, 13 of the 15 phyla identified in the gut and oral microbiota were significantly different between the two sites (after p-value adjustment). Of the top 10 differentially abundant phyla, Bacteroidetes, Verrucomicrobia, Cyanobacteria, Tenericutes, and Euryarchaeota were more abundant in the gut microbiota whereas Proteobacteria, Firmicutes, Actinobacteria, Fusobacteria, and TM7 were more abundant in the oral microbiota (**Supplementary Figure 5A**). At the family level, 55 of 77 identified families were differentially abundant in the gut vs oral microbiota. Of the top 10 differentially abundant families, Bacteroidaceae, Ruminococcaceae, Lachnospiraceae, and Verrucomicrobiaceae were enriched in the gut samples while Streptococcaceae, Pasteurellaceae, Veillonellaceae, Gemellaceae, and Neisseriaceae were enriched in the oral samples (**Supplementary Figure 5B**). At the genus level, 102 of 143 identified genera were differentially abundant in the gut vs oral microbiota. Of the top 10 differentially abundant genera, *Bacteroides*, *Ruminococcus*, *Akkermansia*, and *Lachnospira* were enriched in the gut and *Streptococcus*, *Haemophilus*, *Neisseria*, *Veillonella*, *Selenomonas*, and *Granulicatella* were enriched in the oral samples (**Supplementary Figure 5C**).

We next explored the similarity between the gut and oral microbiota within each cohort. Venn diagrams illustrate that in the BC group, 118 (43.4%) taxa were predominantly fecal, 74 (27.2%) were predominantly oral, and 80 (29.4%) overlapped between the gut and oral samples (**Suppl Figure 6A**). In the DCIS group, 122 (52.1%) taxa were predominantly fecal, 79 (33.8%) were predominantly oral, and 85 (36.3%) overlapped between the two sites (**Suppl Figure 6B**). Finally, of the healthy samples 103 (44.2%) taxa were predominantly fecal, 68 (29.2%) were predominantly oral, and 62 (26.6%) overlapped (**Suppl Figure 6C**).

## DISCUSSION

In this study, we examined the oral and gut microbiomes of women with breast cancer, women with DCIS, and healthy women. Microbiota were analyzed in terms of diversity, taxonomic profiles, and predicted functions. We identified various compositional and functional features of the gut microbiota that differed between women with breast cancer, women with DCIS, and healthy women. In contrast, very few features of the oral microbiota differed between these cohorts.

We found that fecal microbial α diversity, as estimated by the Simpson and Shannon indices, was significantly lower in individuals with BC when compared to healthy individuals, whereas there was no statistically significant difference comparing women with DCIS and healthy women. Reduced α diversity of gut microbiota in women with BC has been reported in several previous studies.^26–30^

Comparing the gut microbiota among the three cohorts, we found that the phylum Firmicutes showed a progressive decrease while the phylum Bacteroidetes showed a progressive increase from healthy to DCIS to BC. This was reflected in the Firmicutes/Bacteroidetes ratio which was significantly lower in the BC group compared to the healthy group. Consistent with the lower F/B ratio in the BC cohort, we also observed significantly greater abundance of the family Bacteroidaceae and genus *Bacteroides* in the BC group vs. the healthy cohort. A higher abundance of the genus *Bacteroides* in BC compared to healthy controls has been reported in previous studies.^27,28,31^ It has been observed that breast cancer risk increases with increasing levels of circulating parent estrogens and decreases with increasing ratios of estrogen metabolites to parent estrogens.^24,25,42,43^ An association has also been observed between urinary estrogens and estrogen metabolites, and the diversity and composition of the fecal microbiome. In postmenopausal women, high fecal microbial diversity is statistically significantly associated with a high ratio of estrogen metabolites to parent estrogens, while the abundance of *Bacteroides* is inversely associated with this ratio.^25^ These data suggest that women with high intestinal microbial diversity are at a lower risk for breast cancer, while women with a high intestinal abundance of *Bacteroides* may be at a higher risk for breast cancer. Our findings are consistent with this.

Specific microbial taxonomic differences vary widely across studies and a definitive disease-associated community structure has not been identified. This may be due to the large variation in gut microbial community composition among humans and/or to technical differences among the studies.

However, several of the differentially abundant taxa in the gut microbiota we observed in our study are consistent with previous studies. In addition to the genus *Bacteroides* mentioned above, we also found the genera *Eggerthella* and *Peptoniphilis* and the class Actinobacteria enriched in the BC vs healthy cohort, consistent with previous studies.^29,44^ In addition, Lachnospiraceae and *Coprococcus* were more abundant in the healthy cohort compared to BC, similar to what was previously reported.^30,44^ Although there are no previous reports on the gut microbiome in women with DCIS, our findings that *Faecalibacterium* was more abundant in healthy vs DCIS gut microbiota and that *Peptoniphilis* and *Actinomyces* were enriched in the DCIS cohort are consistent with previous studies comparing BC with healthy controls.^26,28,44^

Since community structure may be more informative than abundance differences of individual taxa, we clustered the fecal microbiota into guilds/enterotypes. Two clustering algorithms previously used to identify gut enterotypes (ET) are partitioning around medoids (PAMs) or Dirichlet multinomial mixture models (DMMs). Previous studies have suggested that the human gut microbiome can be assigned to no ETs, two ETs (*Bacteroides* and *Prevotella* dominated), three ETs (enriched in *Bacteroides*, *Prevotella*, and *Ruminococcus*), or four ETs (same as the three ETs with an additional *Bacteroides*-dominated cluster).^45–48^ Recently, non-negative matrix factorization (NMF) was used to divide the human gut microbiome into five “enterosignatures” (ES) dominated by *Bacteroides*, Firmicutes, *Prevotella*, *Bifidobacterium*, or *Escherichia.*^49^ The *Bacteroides*, Firmicutes, and *Prevotella* ESs were dominant in adult samples, while the *Bifidobacterium* and *Escherichia* ESs were more often observed in the infant samples. We applied NMF to our fecal microbiota samples and identified 5 guilds. The G5-*Bacteroides* and G1-*Prevotella* guilds were consistent with the previously described ETs/ESs while the G4-Clostridiales guild was similar to the *Ruminococcus* and Firmicutes ETs/ESs.

We found that the G5-*Bacteroides* and G2-Enterobacteriaceae guilds were significantly enriched in the BC samples compared to healthy controls and the G4-Clostridiales guild was significantly enriched in the healthy samples compared to the BC group. Zhu and coworkers,^50^ using PAM clustering, identified 2 enterotypes (*Bacteroides* and *Prevotella* dominated) in fecal samples from women with breast cancer and healthy controls. However, they found no significant relationship between enterotype and disease status. The association of the G5-*Bacteroides* guild with breast cancer is consistent with our results and previous studies suggesting a greater proportion of the genus *Bacteroides* in BC compared to healthy controls.^27,28,31^ The G4-Clostridiales guild was more abundant in the healthy group compared to the BC group. This guild is characterized by a high proportion of the order Clostridiales, the families Lachnospiraceae and Ruminococcaceae, and the genera *Blautia*, *Faecalibacterium*, and *Ruminococcus*. Similar results have been reported in two other studies, where healthy controls had a greater proportion of Clostridiales, Ruminococcaceae, and *Faecalibacterium.*^26,28^

Breast cancer-associated alterations in the gut microbiota likely translate into alterations in gut microbial functions. We utilized Tax4Fun to predict functional pathways from the fecal 16S rRNA sequence data. Pathways enriched in the BC cohort compared to the healthy group included lipopolysaccharide biosynthesis, glycan biosynthesis and metabolism, lipid metabolism, galactose metabolism, and sphingolipid metabolism. Lipopolysaccharide is a potent trigger of systemic inflammation^51^ and has been suggested to play an important role in promoting tumor associated inflammation.^52–54^ Sphingolipid metabolism also plays a role in intestinal inflammation.^55^ Enrichment of these pathways may induce systemic low-grade inflammation, thereby increasing the risk of developing or perhaps exacerbating breast cancer. We do not know if the dysbiosis observed in the BC cohort was present in these women prior to developing breast cancer. Interestingly, we only observed 3 differentially enriched functional pathways between the DCIS and healthy cohorts and of the 16 pathways that differentiated DCIS and BC, most also differentiated BC and healthy gut microbiomes, indicating that the functionality of the gut microbiota from women with DCIS was more like that of healthy women than women with BC.

In contrast to our results from the fecal microbiota, we found no differences in alpha or beta diversity in the oral microbiota of women with BC vs. healthy controls. These results are consistent with the two other published reports on the oral microbiome and breast cancer.^37,38^ While one previous report found no differences in relative abundance of individual taxa,^37^ we identified 1 taxon from the family Ruminococcaceae that was at a significantly greater proportion in the BC cohort. We found no predicted metabolic pathways differentially enriched in the oral microbiota between the BC and healthy groups, which is consistent with the lack of differentially proportional taxa in the oral microbiota. Applying NMF to the oral microbiota we identified seven guilds, however, there were no differences in guild memberships between the BC and healthy groups. We also compared the oral microbiota of women with DCIS and healthy women and found no differences in alpha diversity, very few differentially proportional taxa, no predicted pathway differences, and no differences in guild memberships. Overall, these results indicate that the oral microbiota are very similar between healthy women and women with DCIS or invasive breast cancer.

Breast density, as estimated by mammography, is a strong risk factor for breast cancer. In this study we found that alpha diversity of fecal and oral microbiota did not differ between women with high versus low breast density. Mammographic breast density also was not associated with fecal or oral microbiota beta diversity. In addition, few taxa were differentially associated with high versus low breast density. These results are consistent with a previously published study on mammographic breast density associations with fecal microbiota.^56^

The oral and gut microbiomes are well segregated due to the presence of an oral–gut barrier, represented by their physical distance and the presence of gastric acid and bile.^57–60^ Consistent with previous reports,^6,61,62^ the compositions of the oral and gut microbiota were found to differ greatly in the present study. However, impairment of the oral–gut barrier can allow inter-organ translocation and communication. While inflammatory bowel disease and colon cancer are both conditions where oral taxa are found to be enriched in the gut, recent improvements in quantifying the absolute bacterial load in oral and gut samples has revealed that oral taxa enrichment may be more a reflection of reduced gut bacterial loads.^63^ We analyzed the oral-gut microbiota overlap, and found from Venn diagram analysis that 29%, 36%, and 27% of taxa overlapped across the three cohorts (BC, DCIS, healthy), respectively. These results suggest that oral-gut inter-organ translocation was similar across these cohorts. Intriguingly, we did find that the cariogenic dental pathogen *Streptococcus* was enriched in the gut microbiota of women with breast cancer and DCIS compared to healthy women.

There are several limitations within our study that should be noted when reviewing our results. The first is that we did not control for various factors that are known to affect the microbiome, such as diet, probiotic use, physical activity, and environmental exposures. However, we did control for treatment exposure by collecting samples from the BC and DCIS cohorts prior to any therapy. Second, the sample size of each cohort is small, although still large enough to allow us to identify differences in community composition and individual taxon abundances. Finally, our cohorts were relatively homogeneous with respect to ethnic diversity, limiting our ability to identify microbial features in groups that are disproportionately affected by breast cancer.

In conclusion, our study is the first to analyze and compare both the oral and gut microbiota of women with DCIS, women with BC, and healthy women. Our results suggest that the oral microbiota is unlikely to be an effective risk marker for DCIS or breast cancer. In contrast, there are several distinguishing features in the gut microbiota associated with BC and DCIS and studies might now be designed to deepen our understanding of these associations. In particular, guild-based analysis may provide an ecologically meaningful approach for understanding the relationship between the gut microbiota and breast disease. Lower alpha diversity in women with breast cancer suggests that there is room for interventions targeting the microbiome to improve treatment outcomes and long-term prognosis. Longitudinal studies, for example, might focus on the association between the gut microbiome and chemotherapy, or breast cancer mutations such as BRCA. Interventional studies could be designed to explore the manipulation of the microbiome to determine its role in breast cancer prevention or treatment response. Not least, investigation into specific microbes and the elucidation of the metabolic, biochemical, and immunological processes in which they are involved may help us to better understand how the microbiome can influence the initiation and/or progression of breast cancer.

## MATERIALS AND METHODS

### Participant Selection and Enrollment

Participants eligible for enrollment in this cohort study had a recent diagnosis of IDC or DCIS. Women who had started chemotherapy or hormone treatment, or who had a previous diagnosis of breast cancer, were excluded. Healthy women without a diagnosis of breast cancer or a history of breast cancer served as the control group. Participants with IDC or DCIS were identified at the UCSF Breast Care Center while healthy participants were identified via the ATHENA network (www.athenacarenetwork.org). Participants were enrolled and samples collected between May 2015 and Jan 2017. Ethical approval was obtained from the UCSF Institutional Review Board, and subjects provided written consent for sample collection and subsequent analyses.

### Sample collection, processing, and sequencing

Kits containing material for collecting saliva and stool swab samples were distributed to participants for self-collection. Participants were asked to collect oral samples immediately upon awakening to avoid any immediate effects of daily activities (e.g. eating and brushing teeth) on the oral microbiome. Samples were collected prior to any systemic therapy (including chemotherapy, hormone therapy, and radiation) to avoid therapy-associated effects on the gut and oral microbiomes. Sample swabs were placed in a vial containing a lysis and stabilization buffer that preserves the DNA for transport at ambient temperatures and were stored at - 80°C prior to batch DNA extraction. Samples were lysed using bead beating, and DNA was extracted by a guanidine thiocyanate silica column-based purification method using a liquid-handling robot.^64,65^ PCR amplification of the 16S rRNA gene was performed with primers targeting the V4 variable region (515F: GTGCCAGCMGCCGCGGTAA and 806R: GGACTACHVGGGTWTC TAAT).^66^ In addition, the primers contained Illumina tags and barcodes. Samples were barcoded with a unique combination of forward and reverse indexes allowing for simultaneous processing of multiple samples. PCR products were pooled, column-purified, and size-selected through microfluidic DNA fractionation.^67^ Consolidated libraries were quantified by quantitative real-time PCR using the Kapa Bio-Rad iCycler qPCR kit on a BioRad MyiQ before loading into the sequencer. Sequencing was performed in a pair-end modality on the Illumina NextSeq 500 platform rendering 2 x 150 bp pair-end sequences.

### Data analysis

For the 16S rRNA analysis, the raw reads were joined using join_paired_ends.py script followed by quality-filtering and demultiplexing using split_libraries_fastq.py script in QIIME 1.9.1. The final set of demultiplexed sequences was then selected for Amplicon Exact Sequence Variant (ESV) picking using the DeBlur pipeline. In the pipeline, *de novo* chimeras and artifacts were removed, and ESVs present in less than 10 samples were removed using the *phyloseq* package. The final BIOM file comprising of unique 12,931 ESVs with average 35,378 reads per sample was then used for further analyses.

We used R software packages *phyloseq*^68^ and *microeco*^69^ to conduct analyses of the gut and oral microbiomes. All plots were generated in R using *phyloseq*, *microeco*, *ggplot2*, and *ggpubr* packages. Shannon and Simpson indices were used to estimate α diversity and the variation between groups (β diversity) was tested using permutational multivariate analysis of variance (PERMANOVA). Principle Coordinate Analysis (PCoA) plots based on Bray-Curtis distance at the genus level were employed to illustrate β {Citation}diversity variations.

Key taxa responsible for the differences in the oral and gut microbiotas between the DCIS, breast cancer and healthy cohorts were identified using the LEfSe (Linear discriminant analysis Effect Size) algorithm for biomarker discovery.^70^ We defined a significant α of 0.05 and an effect size threshold of 3 for these analyses. DESeq2 analyses (as implemented in the *microeco* R package) were also used to identify differentially abundant bacterial genera between the groups based on an FDR adjusted p-value cut-off of 0.05.

Functional predictions of the gut and oral microbial communities were performed using Tax4Fun^39^ as implemented in the *microeco* R package, using the SILVA123 database. LEfSe analyses were used to analyze the differences of KEGG pathways between the three cohorts, with a significant α of 0.05 and an effect size threshold of 2.5.

### Guild analyses

Non-negative matrix factorization (NMF) was used to identify bacterial communities or guilds in the gut and oral microbiota samples. NMF was implemented using the *NMF* package in R.^71^ An optimal rank (number of guilds) was determined by performing a rank survey analysis on relative abundance matrices of taxa with ranks = 2 to 15. Unit Invariant Knee (UIK) methodology, implemented with the R package *inflection*^72^ was used to identify the knee/elbow point (optimal rank) of the NMF rank survey plot of residual sum of squares (rss).^73^ NMF analyses were then run using the optimal rank values and the resulting guilds were named according to the most notable taxa in each. The relative abundance of guilds in each sample was determined and the sample was assigned to the guild with the highest relative abundance. The relative abundances of each guild in the three cohorts were compared using Wilcoxon tests.

## Data availability

All fastq files for 16S sequencing were deposited in the NCBI Sequence Read Archive (SRA) (accession number PRJNA1136994). Participant metadata and taxa counts for the gut and oral samples are available for download from https://doi.org/10.5281/zenodo.12775458. A STORMS (Strengthening The Organization and Reporting of Microbiome Studies) checklist ^74^ is available at https://doi.org/10.5281/zenodo.12775468.

## ACKNOWLEDGMENTS

We would like to thank the women who participated in this study. We also thank Dr. Katrine Whiteson for her feedback on the manuscript. This work was supported by the Breast Cancer Research Foundation and the California Breast Cancer Research Program. While engaged in this study, no authors had conflicts of interest to report.

## SUPPLEMENTAL FIGURES

**Supplemental Figure 1.**
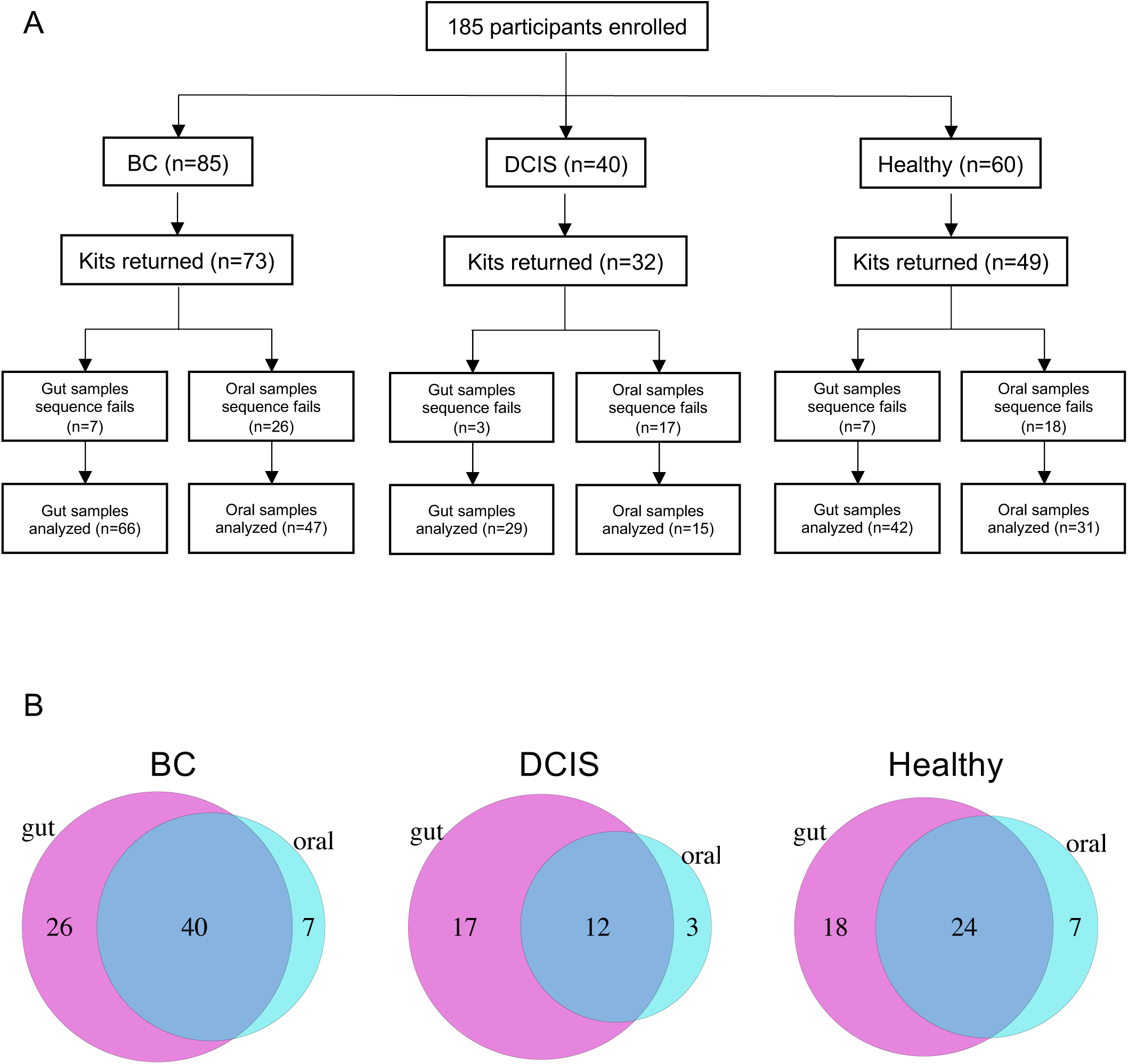
(A) Consort diagram showing participant cohorts and flow of samples in the study. (B) Venn diagrams illustrating the numbers of gut (magenta) and oral (light blue) samples analyzed from women with breast cancer (BC), women with DCIS, and healthy women. Overlapping regions indicate number of participants from which there were paired gut and oral samples.

**Supplemental Figure 2.**
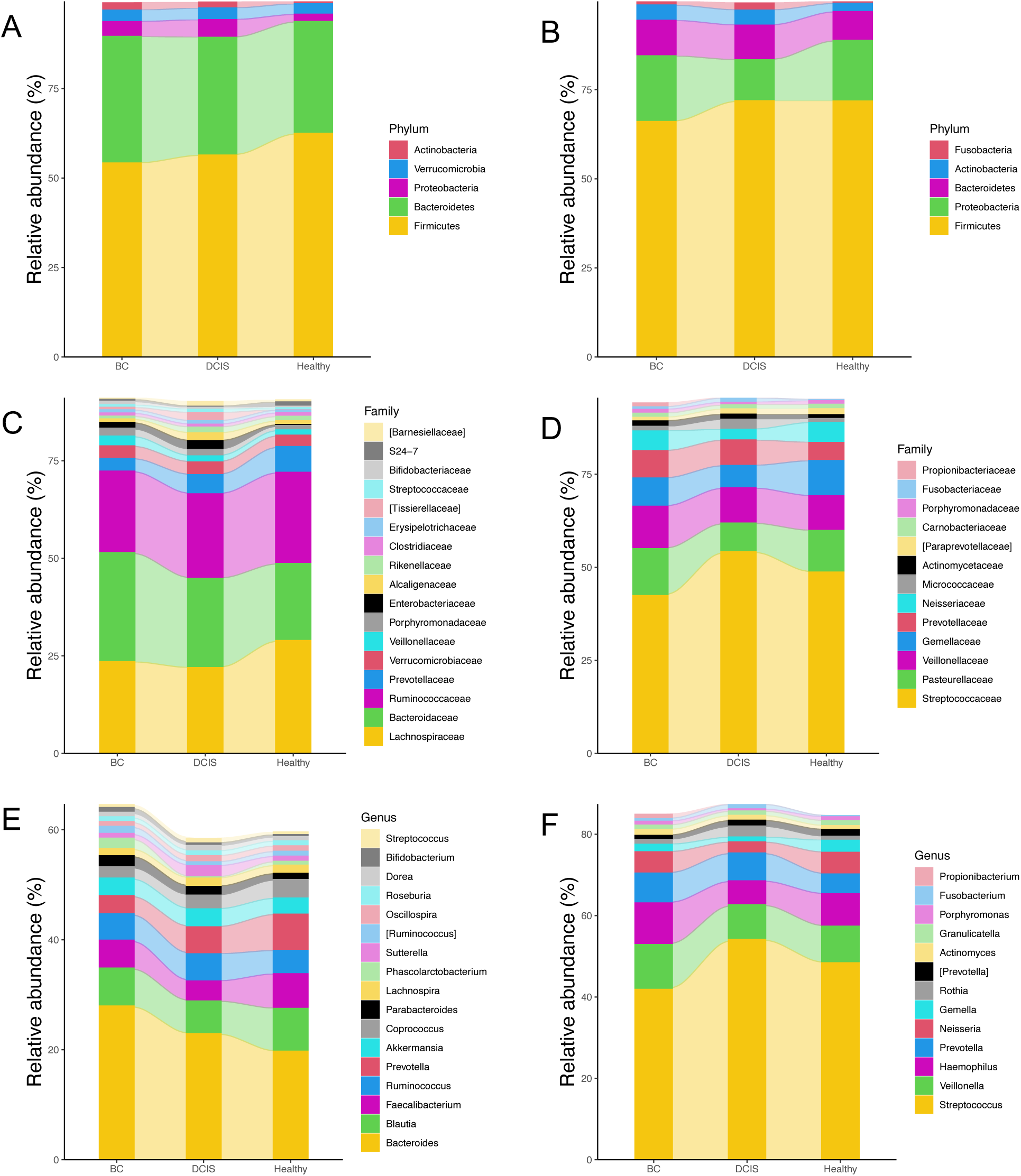
Bar charts of relative abundance of bacteria phyla (A, B), families (C, D), and genera (E, F) in gut (A, C, E) and oral (B, D, F) samples. Only taxa with ≥ 1% relative abundance are shown.

**Supplemental Figure 3.**
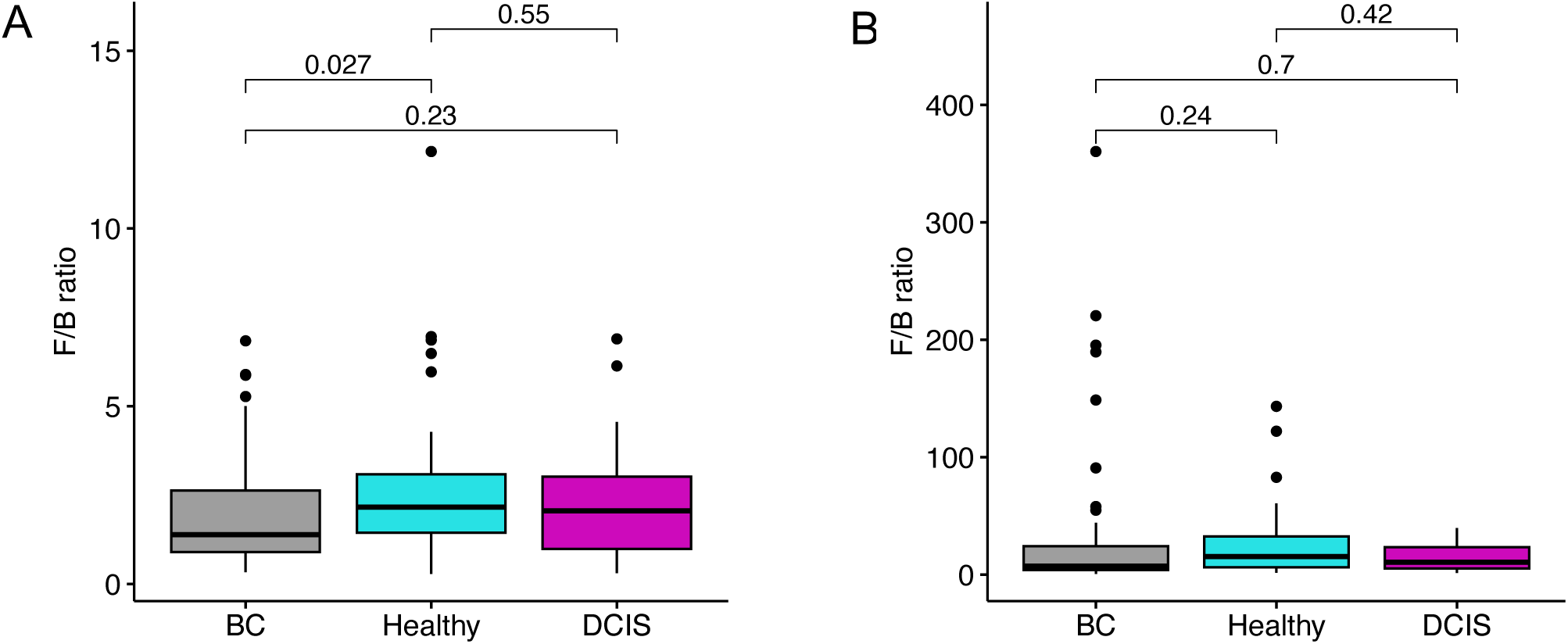
Comparison of the Firmicutes/Bacteroidetes ratio between women with BC, women with DCIS, and healthy women. (A) gut microbiota; (B) oral microbiota. p-values calculated using Wilcoxon tests.

**Supplemental Figure 4.**
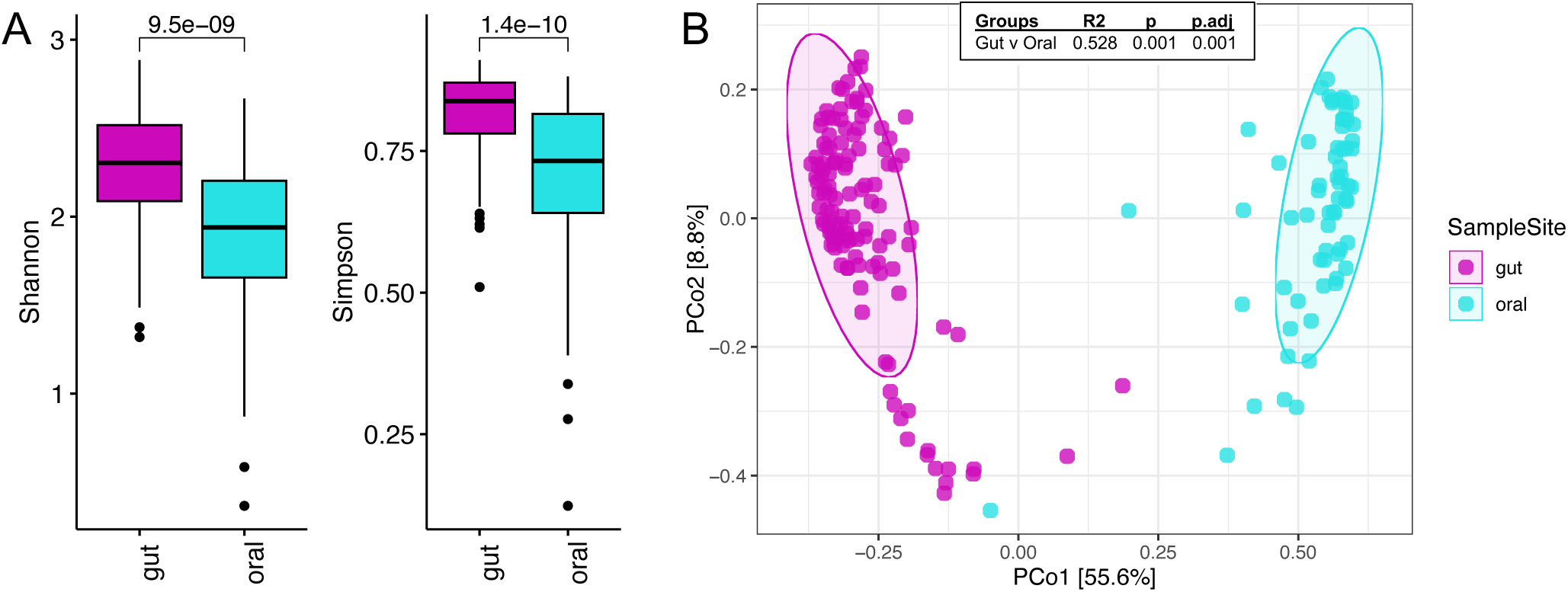
Diversity comparisons of gut vs. oral microbiota. (A) Shannon and Simpson alpha-diversity analyses at the genus level. p-values calculated using Wilcoxon tests. (B) Principal-coordinate analysis (PCoA) based on Bray-Curtis distance at the genus level. R2 and p-values calculated using pairwise PERMANOVA.

**Supplemental Figure 5.**
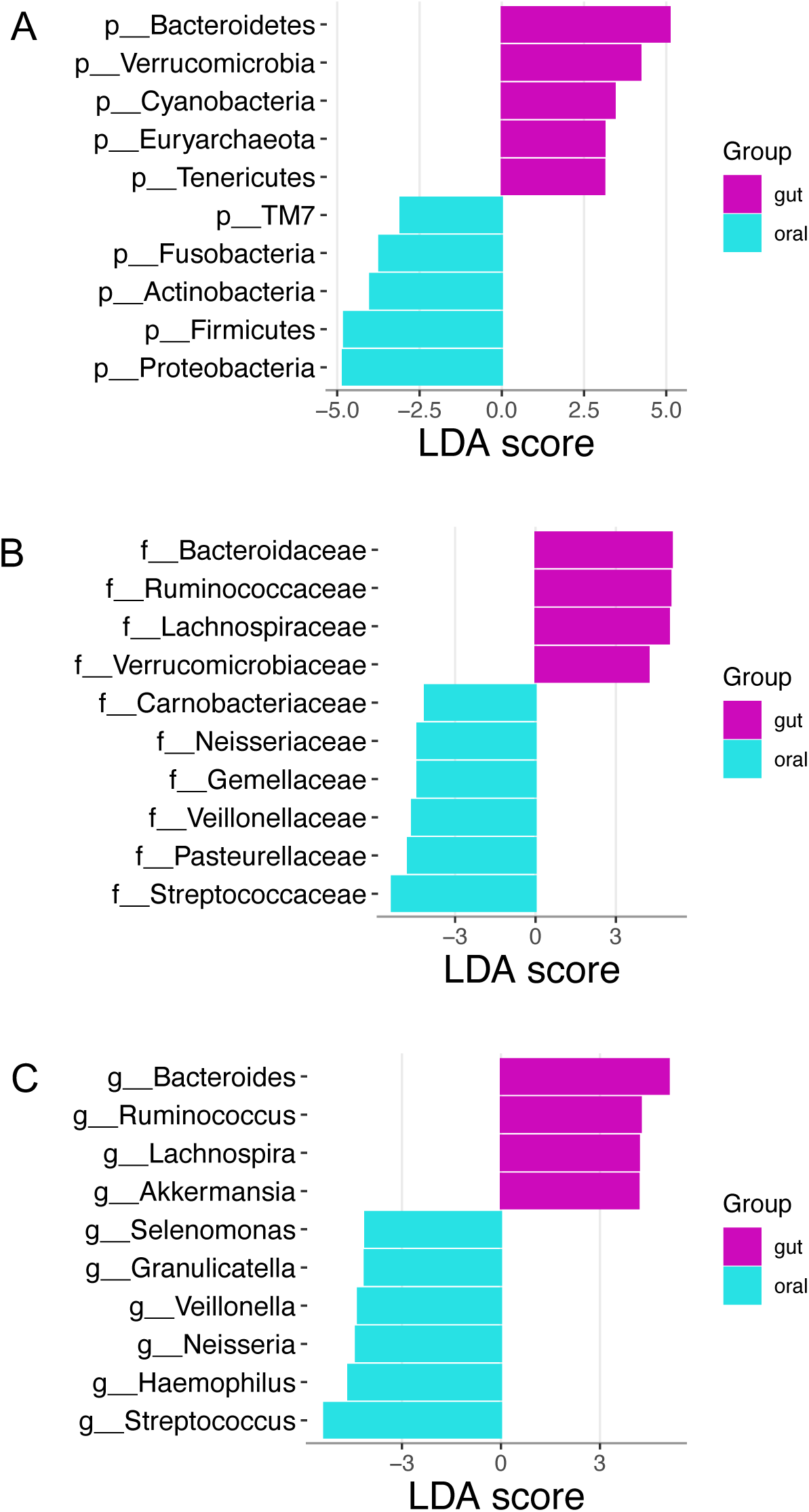
LEfSe analysis comparing gut and oral microbiota at the phylum (A), family (B), and genus (C) levels. The top 10 differentially abundant taxa, based on LDA score, are shown.

**Supplemental Figure 6.**
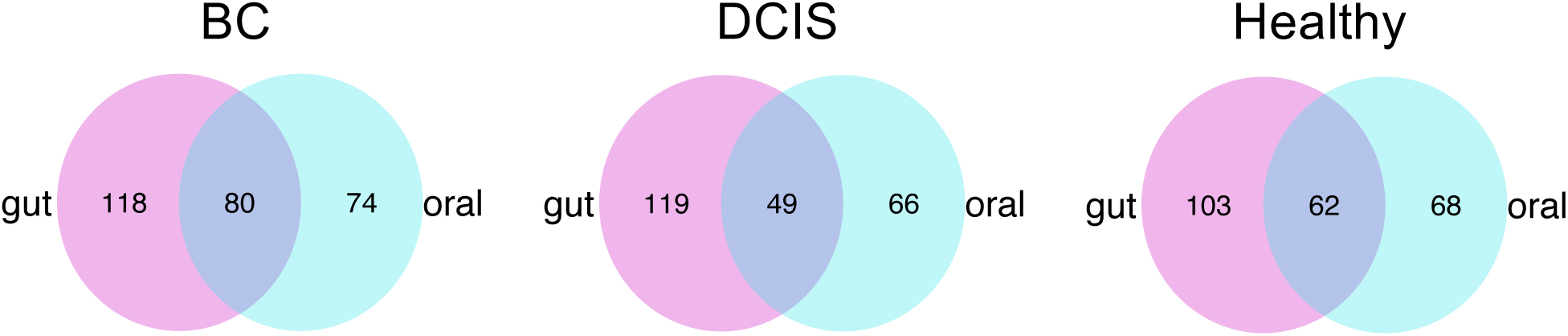
Venn diagrams illustrating the number of shared taxa between gut and oral microbiota in each cohort.

